# Flagellin nebulization enhances respiratory immune responses in the porcine model

**DOI:** 10.1101/2025.03.17.643810

**Authors:** Mara Baldry, Ignacio Caballero, Lisette Ruuls, Delphine Cayet, Yasmine Zeroual, Charlotte Costa, Delphine Beury, David Hot, Yves Le Vern, Nathalie Heuzé-Vourc’h, Ronan MacLoughlin, Arndt G. Benecke, Norbert Stockhofe-Zurwieden, Jean-Claude Sirard

**Author notes:** Equal contribution (Mara Baldry, Ignacio Caballero, Lisette Ruuls, Arndt G. Benecke, Norbert Stockhofe-Zurwieden, Jean-Claude Sirard).

## Abstract

Respiratory delivery of the Toll-like receptor 5 agonist FLAMOD, a recombinant flagellin, offers a promising approach for treating bacterial pneumonia. FLAMOD stimulates the airway epithelium, mobilizing and activating immune cells and effectors to combat infections. While previous evidences were obtained in mouse models, this study represents the first comprehensive assessment of FLAMOD delivered by nebulization in pigs. Our results demonstrate that a single nebulization of FLAMOD did not cause any adverse effects on clinical parameters. Histological analysis supported that FLAMOD treatment led to immune cell infiltration in the lung tissue, indicative of an active immune response. Flow cytometry confirmed granulocyte recruitment in conducting airways. RNA sequencing established immune activation across the respiratory tract, from the nose, trachea, bronchi to the lungs, highlighting innate immunity, bacterial defense, cytokine and chemokine signaling, and granulocyte chemotaxis as key biological pathways. These findings demonstrated the capacity of FLAMOD to induce a robust and common immune response throughout the porcine respiratory system as well as specific compartmentalized immune signatures. This study establishes FLAMOD as a potent activator of innate immunity, providing a proof-of-concept for inhalation-based therapeutic strategies to combat bacterial pneumonia in the clinical setting.

## Introduction

Respiratory tract infections of bacterial or mixed bacterial and viral origin are a major cause of death worldwide (1, 2). This is particularly worrying considering the rise in antimicrobial resistance leading to treatment failure (3). Alternative treatments are thus needed to overcome the burden of antibiotic-recalcitrant bacterial pneumonia (4–6). Stimulation of innate immunity in the respiratory tract is an attractive approach for rapidly and locally activating the host’s defenses against a broad spectrum of microorganisms (7). In mice, prophylactic intranasal administration of the Toll-like receptor 5 (TLR5) agonist flagellin, protects against pneumonia caused by various bacteria (8–11). Moreover, compared with standalone treatments with flagellin or antibiotic, the combination of antibiotic and flagellin synergizes, thus resulting in a lower bacterial load in the lungs and greater protection against *S. pneumoniae* dissemination (12–16). Even within a model of infection with pneumococcus that is resistant to multiple antibiotics in the context of influenza superinfection, the combination of amoxicillin and flagellin leads to bacterial clearance of a 100-fold magnitude higher than that seen in the standalone treatments (13, 14). The prophylactic and protective effects of flagellin are attributed to the recognition by TLR5 on structural cells. After respiratory administration, flagellin predominantly activates airway epithelial cells, triggering the release of cytokines and chemokines that recruit phagocytes, including neutrophils, to the mucosa, where they enhance pathogen clearance (15, 17–20). Concurrently, expression of antimicrobial peptides is induced that can target and neutralize invading pathogens (11). This dual action of cell recruitment and microbicidal activity strengthens mucosal defenses. The selective boosting of airway innate immunity is therefore a conceptually advantageous approach for improving the effectiveness of antibiotic treatment and fighting antibiotic-resistant bacteria (21, 22).

Most studies have been limited to mice, and to inhalation via intranasal administration. However, intranasal administration is not adapted to target the lung tissue in large mammals. For translational purposes towards human use, a more appropriate model to use for respiratory administration of flagellin would be that of the pig. The respiratory tract of pigs (*Sus scrofa domesticus*) is considered similar to that of humans due to anatomical, physiological, and immunological characteristics (23). For instance, the lung lobation, bronchial tree branching patterns, and alveolar structures are similar. The size and structure of the airways, the respiratory rates, tidal volumes, and pulmonary functions are also quite comparable, which is critical for studying drug delivery and infectious diseases (24). Furthermore, pigs have a similar distribution of immune cells in their lungs and exhibit similar immune responses to respiratory infections, which is valuable for studying infectious agents like influenza virus (24, 25). Recent findings highlighted that flagellin stimulates immune signaling in porcine airway epithelial cells, thereby contributing to protection against bacterial pneumonia (26, 27). Aerosol delivery via nebulization is a highly efficient method for administering biologics in liquid form directly to the lungs (28). This approach has been successfully developed for delivery into porcine airways (29, 30). In this study, we present the first instance of delivering flagellin to naïve, healthy pigs using the Aerogen® Solo mesh nebulizer. The objective was to evaluate the effects of flagellin on clinical outcomes and its immune-stimulatory activity.

## Material and Methods

### Flagellin

The custom-designed flagellin used in the present study, *i*.*e*., FLAMOD (recombinant flagellin FliC_Δ174-400_ harboring one extra amino acid at the N-terminus) derives from *Salmonella enterica* serovar Typhimurium FliC (16, 31). FLAMOD was produced in inclusion bodies in *Escherichia coli* by the Vaccine Development department at Statens Serum Institut, Denmark. The flagellin was purified by filtration and chromatography and resuspended in the diluent buffer: 10 mM sodium phosphate, 145 mM NaCl, polysorbate 80 0.02 % (w/v) pH 6.5. The formulation was defined to maintain the molecular integrity and pharmacological activity of flagellin during mesh-nebulisation by the Aerogen^®^ Solo (Aerogen) (32). The immunostimulatory activity was validated using the HEK-Dual™ hTLR5 cell assay (Invivogen). Endotoxin content was found to be <14 endotoxin units per mg of FLAMOD using the *Limulus* assay.

### Animals and housing

Twelve male six-week-old pigs (*Sus scrofa domestica*, line LRxLW TOPIGS 20, Lelystad) were randomized in an unbiased manner by the animal caretakers. The animals were kept in a HEPA-filtered room allocated to three pens with approximately 1 m^2^ floor space per animal. Pens contained straw as floor bedding and were enriched with toys as playing material. Pigs were provided free access to drinking water, which was supplied by drinking nipples. Commercial feed for pigs in nursery units was provided ad libitum. The experiment started after one week of acclimatization. The experiments were conducted in accordance with the Dutch animal experimental and ethical requirements and the project license application was approved by the Dutch Central Authority for Scientific Procedures on Animals (AVD4010020187205) and the experiment plan was approved by the institute’s Animal Welfare Body (permit number: 2018.D-0042)

### Flagellin administration and sampling

Four animals were allocated to each group: group 1 received a nebulized buffer solution (control) and was monitored for 24 hours, group 2 was exposed to nebulized FLAMOD for 4 hours, and group 3 for 24 hours before sampling. Aerosol was administered at day 0 in the morning as follows: pigs were transferred to a separate room, anaesthetized by intramuscular injection with a mixture of Tiletamine-Zolazepam (Zoletil^®^250, 50 mg/ml, Virbac) and Xylazine hydrochloride (Sedaxylan^®^ 20mg/ml, Dechra, NL) at 0.1 ml/kg and then laid into a half pipe in sternal position. Aerosolized treatments were delivered using the Aerogen® Solo nebulizer with the Aerogen® ProX controller system, adapted for pig snouts as previously described. (29, 30). A standard animal mask with a rubber face seal was modified to include a one-way valve, ensuring unidirectional airflow, and filters at both the entry and exit ports to prevent environmental exposure to FLAMOD. The nebulizer, integrated with an Aerogen® 22 mm T-piece, was incorporated into the mask. During inhalation, aerosolized medication was entrained in the inspiratory flow and delivered to the respiratory tract. Exhaled air was directed exclusively through the one-way valve. Previous studies indicated that approximately 30% of the nominal dose reaches the pig lungs (29, 30). FLAMOD (1 ml at 1.5 mg/ml) was administered over 5 minutes, while control animals received 1 ml of buffer. Treatments began with the diluent buffer-treated group to establish a baseline, followed sequentially by the FLAMOD-treated groups. Group 2 pigs were necropsied 4 h post-administration, while groups 1 and 3 were necropsied at 24 h. Blood samples (heparinized blood, serum, and RNA-stabilized blood in Tempus™-Thermofisher) were collected the day before treatment and immediately prior to necropsy. For the clinical assessment, animal weight was recorded one day before and one day after treatment (groups 1 and 3 only) and rectal body temperatures were measured in the morning before treatment and at 4 h and 24 h after treatment; general health observations were conducted twice daily from seven days before treatment until one day after treatment. Specific clinical signs in the upper and lower respiratory tract like coughing, sneezing, nasal/ocular discharge and respiration were also recorded.

### Clinical analysis and biochemistry

Hematological analysis was performed by using an automated hematology analyzer (pocH-100iV, Sysmex) to record white blood cell count, the % of lymphocytes/monocytes and neutrophils, red blood cell counts, hematocrit, hemoglobin and platelet numbers.

### Pathology

At necropsy, tissue specimens were collected from the anterior, middle and caudal lobes of lungs. Tissues were fixed in 4% neutral-buffered formaldehyde, embedded in paraffin, and histological sections of 4 μm thick were stained with hematoxylin and eosin and analyzed semi-quantitively in a blinded manner by a qualified veterinary pathologist (N.S-Z). Histopathological scoring was performed as described previously (33). Briefly, three lung lobes were assessed for inflammatory infiltrates in the bronchial wall, bronchi, peribronchial interstitium, alveolar wall, and alveolar lumen. Lesions were scored on a scale from 0 (no findings) to 3 (extensive manifestation), and scores were averaged.

### RNA extraction

Nasal, tracheal, bronchial, lung tissues and blood samples were collected and preserved in NucleoProtect® RNA (Macherey-Nagel, Duren, Germany) at −80°C. Tissues were thawed and dried on paper towels and any visible cartilage was removed. The remaining tissue was disintegrated with an UltraTurrax homogenizer (IKA-Werke, Staufen, Germany) into 1-3 ml RA1 lysis buffer (Macherey-Nagel) supplemented with 2% TCEP (Sigma) and total RNA extraction was conducted with NucleoSpin^®^ RNA Plus kit (Macherey-Nagel). Lung RNA was eluted in 60 μl RNAse-free water. For nasal, bronchial and tracheal samples, 350 μl of lysate was added to 595 μl water and 10 μl proteinase K (20 mg/ml), and incubated at room temperature for 10 min, and at 55°C for 10 min. After centrifugation, the supernatant was mixed with 490 μl of 100% ethanol and RNA was eluted in 30 μl RNAse-free water. Blood samples (3 ml) that were collected in Tempus™ tubes and stored at −20 °C, were thawed and the stabilized blood was diluted up to 12 ml with D-PBS (Gibco), vortexed for 30 sec, and centrifuged at 4°C for 30 min at 3000 *g*. The pellet was dissolved in 1 ml RA1 buffer + 2% TCEP and 150 μl was used for extraction. Blood RNA was eluted in 80 μl RNAse-free water.

### RNA sequencing and differentially expressed gene (DEG) analysis

RNA quality was evaluated using the TapeStation 4200 (Agilent Technologies), and quantified by spectrophotometry (Nanodrop, Thermo Fisher), and fluorimetry (Qubit, Thermo Fisher). The libraries were prepared using the QIAseq stranded mRNA library kit as well as QIAseq FastSelect-Globin Kit for blood samples (Qiagen), quantified by quantitative PCR (KAPA Library Quantification Kit for Illumina platforms, KapaBiosystems) and pooled in an equimolar manner prior to sequencing on a NovaSeq sequencer (Illumina) in 2 × 150 bp. RNA sequencing data (FASTQ files) were mapped and annotated to the pig genome (Ensembl Sscrofa11.1, GCA_000003025.6) using the STAR alignment software (STAR 2.7.10a; (34)). Genome scaffolds were generated with 149 nucleotide lengths. Differentially expressed gene (DEG) kinetic analysis was performed using the NeONORM method for data normalization (35). Genes (Ensembl IDs) with p-adjusted value < 0.05 (36) were considered as differentially expressed. Metascape was used to identify statistically enriched Gene Ontology (GO) biological processes associated with the various respiratory compartments (37). To leverage advanced human GO annotations, we used Ensembl orthology mappings (homologues and paralogues) to convert *S. scrofa* gene IDs to *Homo sapiens* where possible. Sequencing data have been deposited in the National Center for Biotechnology Information’s Gene Expression Omnibus repository under the accession number GSE289859 and Sequence Read Archive repository under the accession number PRJNA1224715.

### ELISA

Concentration of IL-12/IL23 p40, IL-10 and TNF cytokines in the bronchoalveolar lavage and serum of flagellin-treated pigs and controls was measured using specific ELISA kits (Porcine Quantikine ELISA kits from R&D systems) following manufacturer’s instructions. Standard curves were created for each cytokine and the concentration calculated by fitting a four-parameter logistic curve. Limit of quantification were as follows: IL-12/IL-23p40 (IL-12p40): 47.0 pg/ml; IL-10: 31.3 pg/ml; TNF: 23.4 pg/ml.

### Flow cytometry

Broncho-alveolar lavages (BAL) were collected post-mortem by instilling and re-aspirating two 15-mL aliquots of sterile DMEM high glucose supplemented with 100 IU/ml penicillin, 2.5 μg/ml amphotericin and 50 μg/ml gentamicin. Red blood cells were lysed with an erythrocyte lysis buffer and washed with DMEM supplemented with 10% FBS and 100 IU/ml penicillin before staining of cell surface markers. Cell surface marker stainings were performed in PBS containing 2 mM EDTA supplemented with 5% porcine serum and 5% goat serum for 30 min in ice. Primary antibodies were used as follows: anti-MHCII (mouse IgG2a, clone MSA3, dilution 1/250; Kingfisher Biotech); anti-CD172a (mouse IgG2b, clone 74-22-15a, dilution 1/500; BD Biosciences); anti-CD163 PE-conjugated (mouse IgG1, clone 2A10/11, dilution 1/20; BIO-RAD). Secondary antibodies anti-mouse IgG2a coupled to PE-Cyanine7 and anti-mouse IgG2b couple to APC-Cyanine 7 were purchased from Invitrogen and used at a 1/200 dilution. Cell viability was assessed using the Fixable Viability Dye eFluor™ 450 (ThermoFisher). BAL samples were acquired on a BD LSR Fortessa X-20 (BD Biosciences, San Jose, CA, USA) and data was analyzed using Kaluza analysis software v2.1. Neutrophils and alveolar macrophages were classified as described previously (38), where neutrophils were defined as MHCII^null^/CD163^null^/CD172a^+^, while alveolar macrophages were defined as MHCII^+^/CD163^+^/CD172a^+^.

### Statistical analysis

Results were expressed as individual values and median or mean. Groups were compared using a Kruskal-Wallis one-way ANOVA by controlling the false Discovery rate. Statistical analyses were performed using GraphPad Prism software (version 9.5.1, GraphPad Software Inc., San Diego, CA), and the threshold for statistical significance was set to p<0.05.

## Results

### A single nebulization of FLAMOD is well tolerated with no signs of safety concerns

Using a specifically designed nose mask connected to the Aerogen Solo mesh nebulizer, anesthetized pigs were exposed for 5 min to aerosolized vehicle diluent buffer, (control; terminal sampling at 24 h), or FLAMOD in the diluent buffer with an administered dose at 1.5 mg per animal with terminal sampling at 4 h post-nebulization and 24 h post-nebulization. Under these conditions, approximately 30% of the nominal dose is expected to be delivered to the pig lung (30), and thus, we estimated that each animal had a lung deposition of approximately 0.5 mg FLAMOD. There were no observed adverse effects as measured by body temperature, body weight and respiratory rate (**Figure 1a-c**). Any signs of coughing, sneezing, nasal or ocular discharge, and respiratory distress were not observed. The hematology was also analyzed at 4 h or 24 h post-nebulization to define any systemic adverse effects on the blood compartment (**Figure 1d-j**). There were no significant differences in numbers of red blood cells, platelets, white blood cells, hemoglobin or hematocrit between the control and the FLAMOD-receiving groups post-nebulization. Collectively these results show that nebulization of a single FLAMOD dose is well tolerated and has no safety concerns.

**Figure 1.**
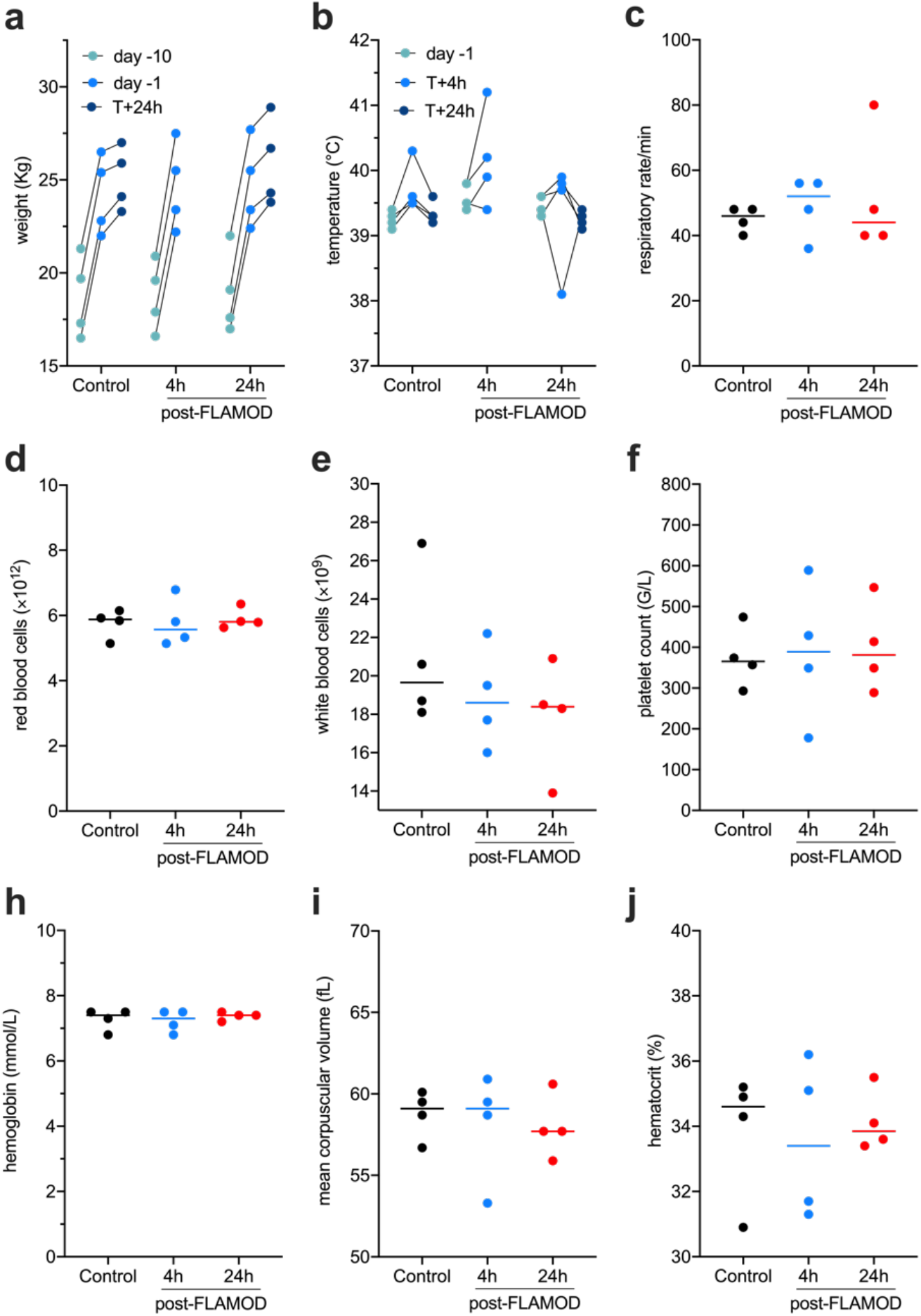
A single nebulization of FLAMOD did not alter body weight, temperature, respiratory rate or hematology counts. Pigs were nebulized with diluent buffer (control group) or FLAMOD at 1.5 mg per pig. Animals were analyzed and samples were collected 4 h post-nebulization (4 h post-FLAMOD group) or 24 h post-nebulization (control and 24 h post-FLAMOD groups). (**a**) Body weight was recorded at day −10, day −1 (pre-nebulization) and at T+24 h post-nebulization. (**b**) Body temperature was measured at day −1, 4 h post-nebulization (T+4h) and 24 h post-nebulization (T+24h). (**c**) Respiratory rate was measured post-nebulization. (**d-j**) Hematological analysis was performed at 4 h and 24 h post-nebulization using an automated hematology analyzer. Statistical analyses using one-way ANOVA did not show any significant differences between groups.

### Nebulization of FLAMOD promotes inflammatory cell mobilization into lungs and triggers local and systemic immune responses

As part of the assessment, both macroscopic and microscopic evaluations of the lungs in FLAMOD-treated pigs were conducted. Macroscopic examination did not reveal any pathological changes (including increased mucosal redness or augmented secretion in the nasal mucosa, nasopharyngeal area, or trachea) in any group. Moreover, lung retraction following exenteration remained unchanged in the FLAMOD-treated groups as discoloration of lung tissue or increased bronchial excretions were not observed upon necropsy. Microscopic histopathological analysis revealed that FLAMOD nebulization led to a progressive increase in inflammatory cell infiltration in both the bronchial and alveolar compartments over time compared to the buffer control group **(Figure 2a-c)**. The histological scoring of tissue sections, including the bronchi, bronchial wall, peribronchial region, alveolar insterstitium, and alveoli, correlated with the observed images and cell infiltrates **(Figure 2d)**.

**Figure 2.**
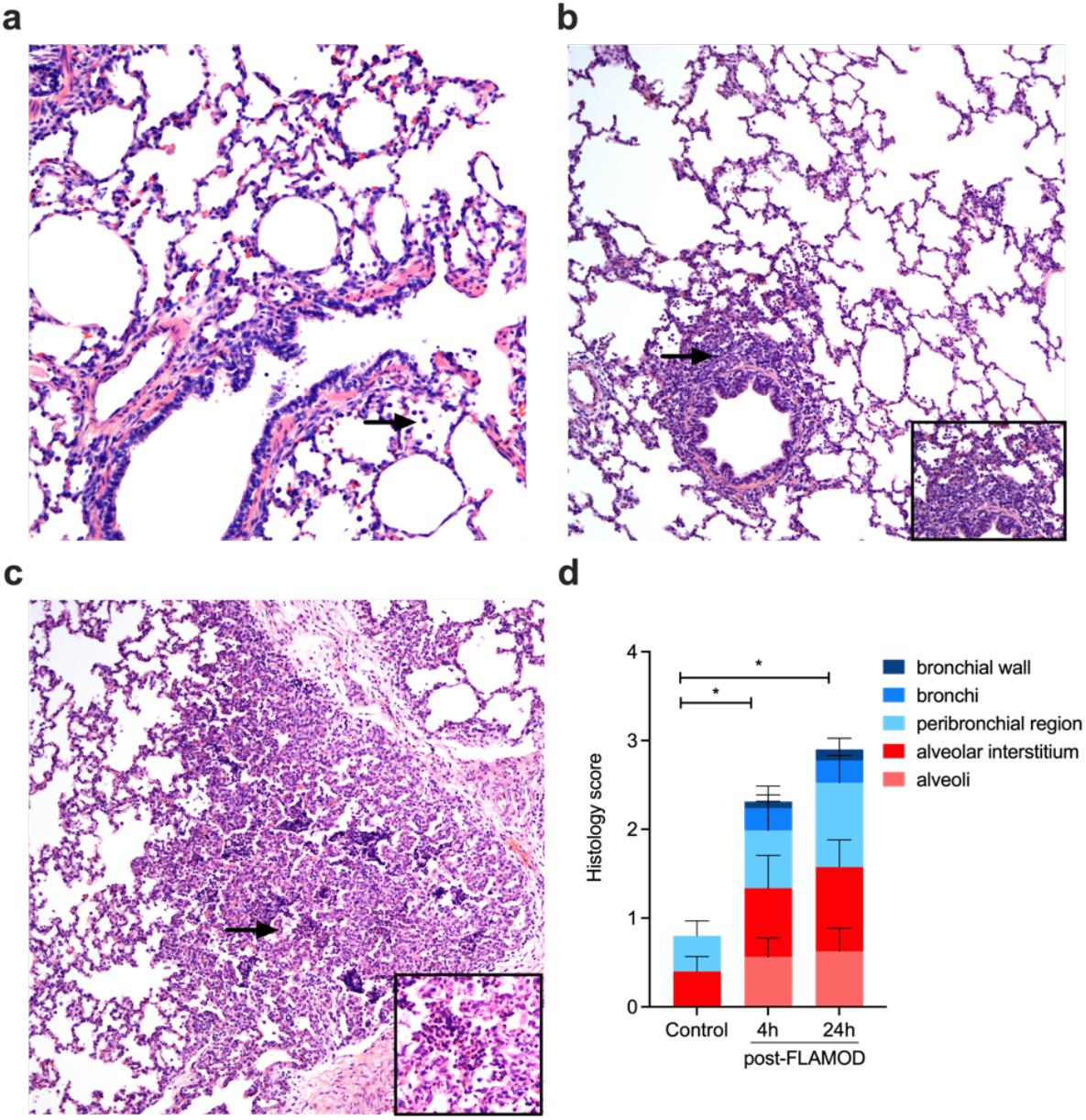
Nebulization of FLAMOD induces immune cell infiltration into the lungs. Pigs were nebulized with diluent buffer (control group) or FLAMOD at 1.5 mg per pig. Animals were sacrificed and lung tissues were sampled 4 h post-nebulization (4 h post-FLAMOD group) or 24 h post-nebulization (control and 24 h post-FLAMOD groups). Lung tissue sections were prepared and stained with hematoxylin-eosin for **(a-c)** microscopy analysis at a 100X magnification and **(d)** histopathological scoring. **(a)** Tissue from control pigs exhibited minimal mononuclear cells (arrow) in the lung alveoli. **(b)** Lungs of pigs 4 h post-FLAMOD showed a mild localized, increase in mononuclear cells and neutrophil infiltration (arrow) in the peribronchial region (insert) and alveoli. **(c)** Lungs of pigs 24 h post-FLAMOD displayed incidentally a focal extensive infiltration of mononuclear cells and neutrophils (arrow and inset), along with interstitial inflammatory cell infiltration in the alveolar septa. **(d)** Histology scoring was performed to quantify inflammatory cell infiltration across different lung lobes on a scale of 0 (no findings) to 3 (extensive infiltration). The cumulative mean scores from three lung lobes are shown. Statistical analysis was performed using one-way ANOVA.

To further characterize the respiratory immune response to FLAMOD, bronchoalveolar lavage (BAL) samples were analyzed by flow cytometry **(Figure 3)**. The results demonstrated that FLAMOD promotes granulocyte recruitment into the conducting airways, as indicated by a significantly increased percentage of BAL neutrophils following nebulization **(Figure 3a)**, accompanied by a decrease in alveolar macrophages **(Figure 3b).** FLAMOD also modulated soluble immune mediators, with IL-12 and TNF levels elevated in BAL fluid at both 4 h and 24 h post-nebulization, while IL-10 production was detectable only at 24 h (**Figure 3c**). In addition, FLAMOD nebulization enhanced IL-10 and IL-12 production in circulating blood (**Supplementary Figure 1**), whereas systemic TNF levels remained unaffected. These findings suggest that delivery of FLAMOD in the airways by nebulization enhances innate immunity by promoting the recruitment of immune cells, particularly neutrophils, from circulation to the lungs and conducting airways.

**Figure 3.**
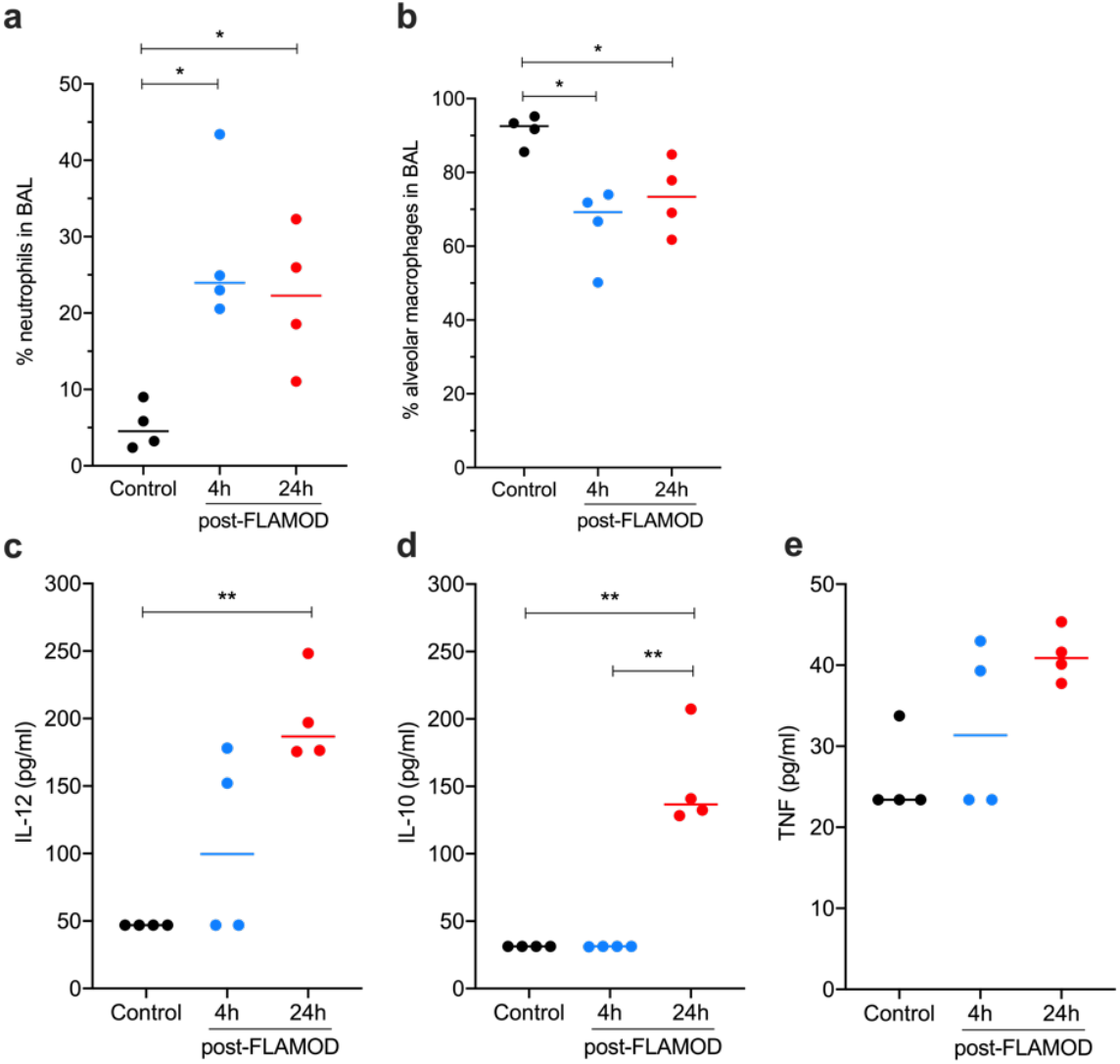
Nebulization of FLAMOD modulates immune cell populations in the lungs and promotes local immune responses. Pigs were nebulized with diluent buffer (control group) or FLAMOD at 1.5 mg per pig. Animals were analyzed 4 h post-nebulization (4 h post-FLAMOD group) or 24 h post-nebulization (control and 24 h post-FLAMOD groups). Bronchoalveolar lavages were sampled for further analysis of cells and cytokines. **(a-b)** Flow cytometry evaluation of (**a**) neutrophil percentage identified as MHCII^null^CD172a^+^CD163^null^, and **(b)** alveolar macrophages identified as MHCII^+^CD172a^+^CD163^+^. **(c-e)** Production of cytokines IL-12 **(c)**, IL-10 **(d)** and TNF **(e)** was evaluated by ELISA. Each symbol represents one animal. Statistical analysis was performed using one-way ANOVA.

### Transcriptional analysis of tissue response shows stimulation of innate immunity in a compartment-dependent manner

Aerosol inhalation via mask nebulization facilitates the delivery of the drug product to multiple regions of the respiratory tract, including the nasal cavity, trachea, bronchi, and the peripheral lung. The observed immune cell infiltration into both the alveolar and bronchial compartments of the lungs provides strong evidence that FLAMOD effectively activates local immune responses (**Figures 2 and 3**). To comprehensively evaluate the potential of FLAMOD to stimulate immune activity across different regions of the respiratory tract, we conducted transcriptional profiling of the nasal mucosa (nose), trachea, bronchi, and lung tissues following administration of either the diluent buffer or FLAMOD. This analysis was performed using RNA sequencing (RNA-seq) **(Figure 4)**.

**Figure 4.**
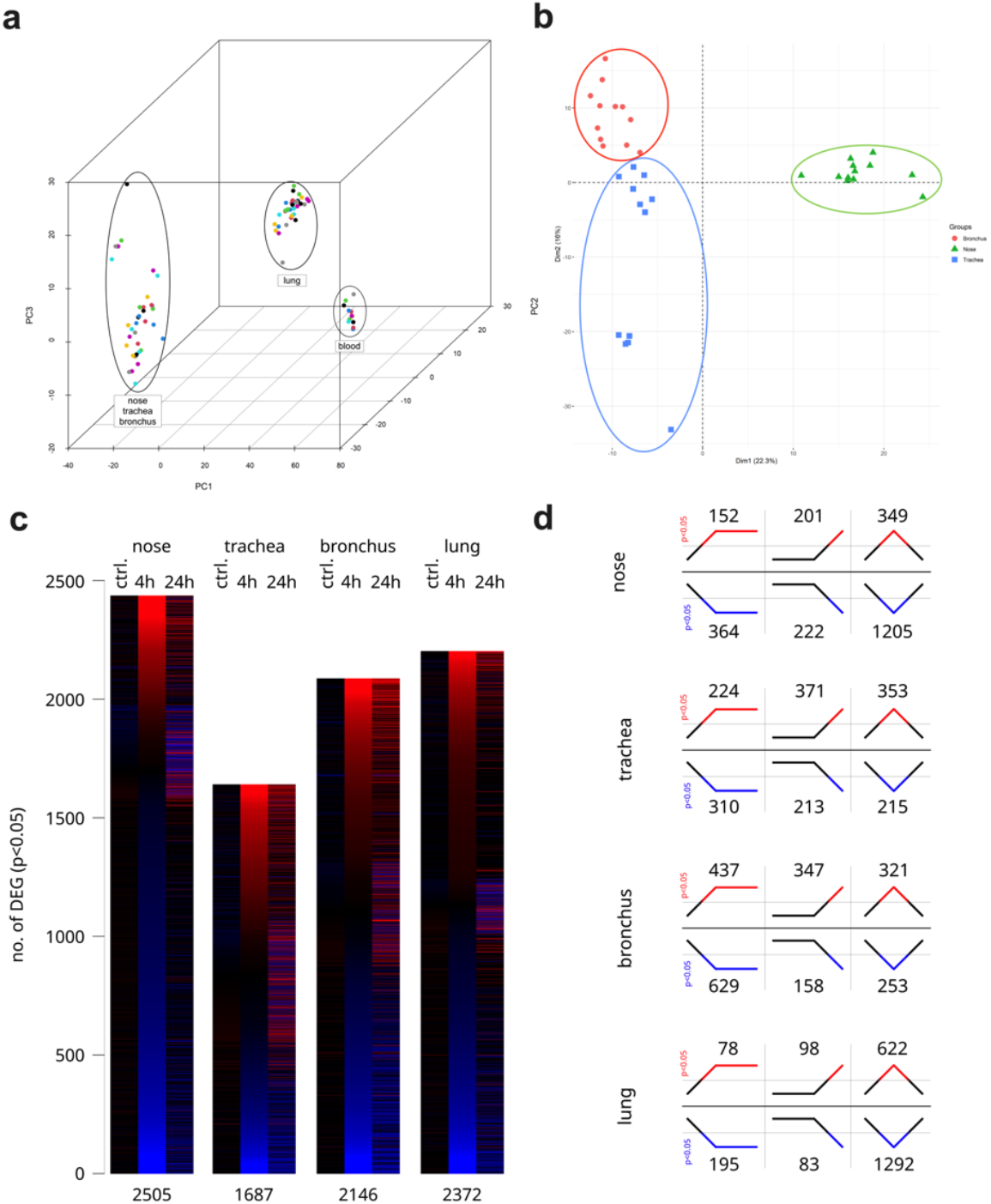
FLAMOD nebulization results in compartmentalized responses. Pigs were nebulized with diluent buffer (control group; Ctrl) or FLAMOD at 1.5 mg per pig. Animals were analyzed 4 h and 24 h after FLAMOD nebulization. Total RNA was extracted and subjected to RNA sequencing. **(a-b)** Tissue classification by PCA plots. **(c)** Heatmaps illustrating the total number and relative expression change of regulated genes (p<0.05) across the different compartments. **(d)** Time series analysis of each compartment showing the patterns of gene regulation as represented by six different line patterns, where the total number of genes statistically matching each pattern are indicated above or below the respective pattern.

Principal component analysis (PCA) was employed to classify the tissues based on their transcriptional profiles. The PCA results clearly segregated the samples into distinct groups corresponding to blood, lung, and the respiratory compartments, including the nose, trachea, and bronchus (**Figure 4a)**. This separation highlighted the unique gene expression patterns associated with each tissue type and the specificity of FLAMOD’s effects. A more refined PCA focusing exclusively on the nose, trachea, and bronchus further distinguished these three compartments (**Figure 4b)**. Additionally, weighted gene co-expression network analysis (WGCNA) identified specific module signatures that also differentiated each respiratory compartment (**Supplementary Figure 2**). RNA-seq analysis revealed significant changes in gene transcription at 4 h and 24 h after FLAMOD nebulization compared to the control group, with the most pronounced transcriptional changes occurring at 4 h across all compartments (**Figure 4c** and **Supplementary file 1**). Noticeable differences were also observed with regards to the total number of differentially expressed genes (DEG) per compartment. Further examination using time-series analysis uncovered distinct temporal expression patterns, highlighting both compartment-specific differences in gene regulation and variations in the number of DEG following each transcriptional trajectory upon FLAMOD administration. These findings suggest that FLAMOD elicits dynamic and regionally specific transcriptional responses throughout the respiratory tract (**Figure 4d**). An integrated analysis of DEG across the nose, trachea, bronchus, and lung was conducted using Metascape (37) (**Supplementary Figure 3a-b**). This analysis revealed that flagellin aerosol delivery elicits a highly coordinated transcriptional response across the respiratory compartments. Analysis of individual respiratory compartments identified significant modulation of biological pathways following FLAMOD nebulization (**Figure 5a**). Several immune-related pathways were commonly enriched across the nasal mucosa, trachea, bronchus, and lung, including responses to bacterial pathogens, inflammatory processes, innate immune activation, and cytokine/chemokine signaling. In addition to immune pathways, commonly regulated processes involved synaptic signaling, transmembrane transport, and cell-junction organization, suggesting broader physiological effects beyond immune activation. An integrated gene ontology analysis identified a diverse array of enriched functional clusters across all respiratory compartments. Notably, inflammatory responses and cilium movement were among the top 20 most significantly enriched clusters (**Supplementary Figure 3c-d**). A detailed examination of the top regulated genes within selected pathways revealed that genes associated with granulocyte responses (namely *CSF3R, CXCL8, CXCR1* and *S100A8*), cytokine/chemokine signaling (*IL12B, IL1A, CXCR12, CXCL6* and *CXCL11*), and antibacterial defense (*PGLYRP2, C1QTNF* family and *CHIT1*) were most strongly upregulated in the trachea and bronchi, followed by the nasal mucosa and lung (**Figure 5b**). Notably, FLAMOD had a distinct impact on cilium organization and motility specifically within the nasal compartment, where it downregulated *BBOF1, CIAP100* and *KIF27* which are key genes involved in these processes (**Figure 5a-b**), thus potentially influencing mucociliary clearance. To further validate the transcriptional changes identified by RNA-seq, a subset of differentially expressed immune-related genes was confirmed using RT-qPCR across multiple respiratory compartments (**Supplementary Figure 4**). Blood was also analyzed with RNA-seq. A transcriptional pattern similar to that in respiratory compartments was observed in the blood of the nebulized pigs (**Supplementary Figure 5**). Collectively these findings demonstrate that while the magnitude and patterns of gene regulation vary by compartment, FLAMOD nebulization in pigs robustly activates gene expression, particularly in pathways related to immune responses, across all exposed regions of the respiratory tract.

**Figure 5.**
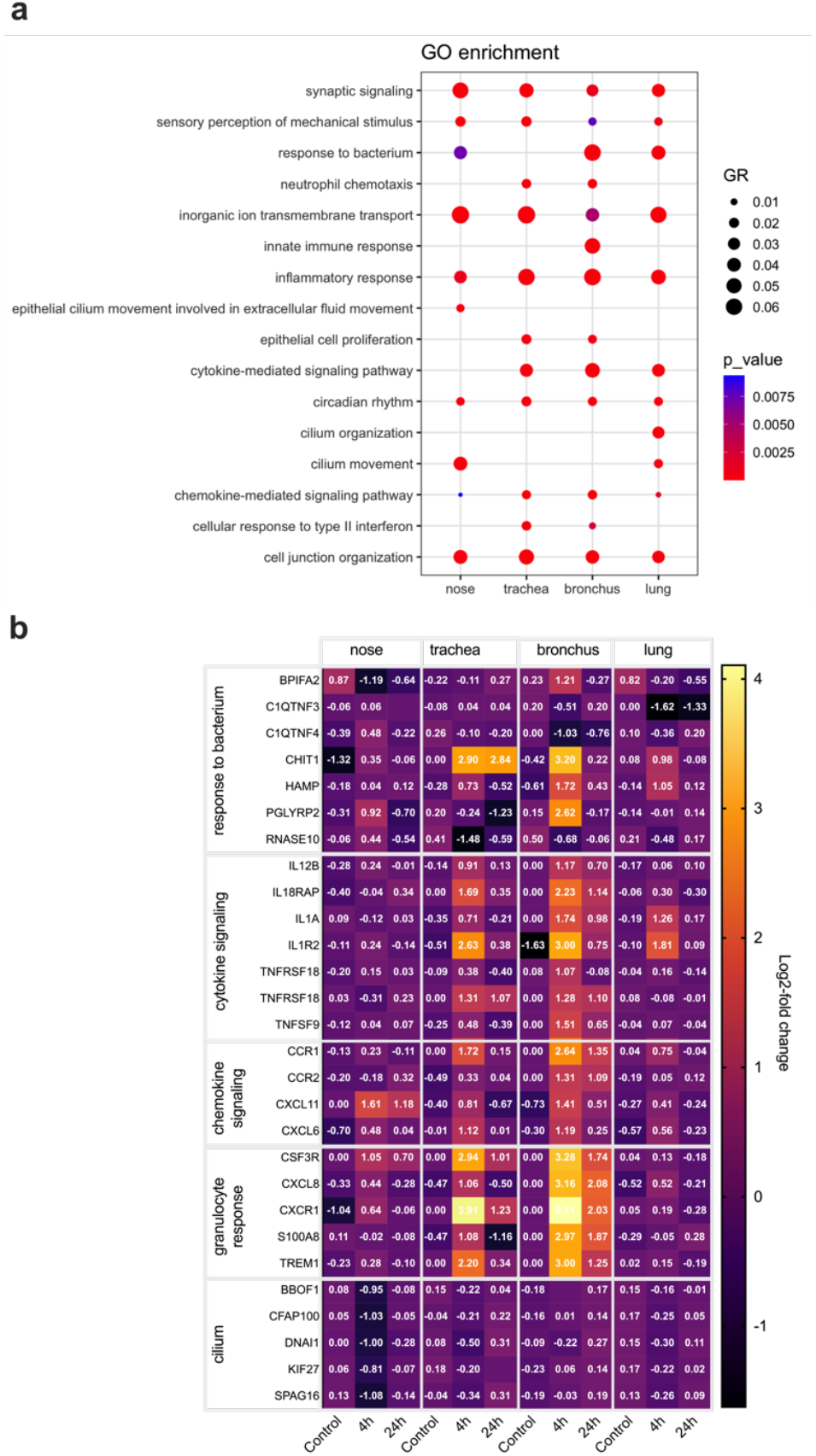
Immune pathways are modulated in respiratory tissues by inhaled FLAMOD. Pigs were nebulized with diluent buffer (control group) or FLAMOD at 1.5 mg per pig. Animals were analyzed 4 h and 24 h after FLAMOD nebulization. Total RNA was extracted and subjected to RNA sequencing. **(a)** Immune pathways across compartments were identified using Metascape. The common biological processes enriched across the different compartments are represented. The gene ratio (GR) and p_value scales indicate the number of genes enriched for in each pathway and the significance of enrichment, respectively. **(b)** Heatmaps of top regulated genes within selected pathways. Annotated values per cell indicate log_2_(fold change).

## Discussion

In this study, we provide the first evidence of aerosolized delivery of the TLR5 agonist FLAMOD in pigs using a vibrating mesh nebulizer. Our findings demonstrate that a single nebulization of FLAMOD is well tolerated over 24 h and induces robust immune activation throughout the respiratory tract, including the nasal mucosa, trachea, bronchi, and lungs. This immune response is characterized by the production of pro-inflammatory mediators in both the conducting airways and circulating blood, as well as the infiltration of immune cells - particularly neutrophils—into the lungs and airways. Transcriptomic analysis revealed that FLAMOD triggers transient but significant changes in gene expression, with strong upregulation of pathways involved in innate immunity and antimicrobial defenses. These findings reinforce the proof-of-concept that mucosal TLR5 activation can enhance the host’s ability to combat respiratory infections, positioning FLAMOD as a promising immunomodulatory strategy for pneumonia prevention and treatment.

Consistent with prior studies in mice(12–16), our findings confirmed that the TLR5 signaling mediated by nebulized FLAMOD in the respiratory tract, induced neutrophil infiltration in porcine lung tissue. The ability to replicate this pharmacodynamic effect in pigs—a species with strong physiological and immunological similarities to humans—supports FLAMOD’s potential for broader clinical application. These results evidenced the strong evolutionary conservation of TLR5 across species (39).

Nebulization has been widely used in porcine models to study pulmonary drug delivery, including the administration of surfactants in acute respiratory distress syndrome models (40) and nebulized antimicrobials like bacteriophages and colistin in ventilator-associated pneumonia (41, 42). Nebulization studies demonstrated superior lung deposition with nebulization over intravenous administration (42). Here we show that nebulization of FLAMOD delivers the biologic to the airways of pigs in a safe and effective manner. Although vibrating mesh nebulizers ensure even distribution across the respiratory tract, deposition rates are lower compared to intranasal or mucosal atomization (30). Previous murine studies indicate that lung immune responses are elicited with 1–2.5 μg of flagellin (13, 14), while porcine airway epithelial cells respond maximally at 100 ng/mL (27). Thus, respiratory immune activation in pigs receiving an estimated lung-deposited dose of 500 μg of FLAMOD—substantially higher than previously tested doses—demonstrates robust efficacy without observable adverse effects. This finding highlights the strong local tolerance to FLAMOD inhalation and supports its favorable safety profile for airway-targeted delivery.

Our study demonstrates FLAMOD’s “hit and run” mechanism, wherein transient gene modulation occurs from the nasal passages to the bronchi and conducting airways. Peak transcriptional activation at 4 hours post-nebulization was followed by partial resolution at 24 hours. Enriched pathways included inflammatory responses, and chemokine signaling, aligning with known flagellin-induced innate immune activation in mice (10, 14, 16). Chemokines such as CCL20, CXCL2, and CXCL8 (IL-8), known to mediate neutrophil chemotaxis were significantly upregulated, corroborating the enrichment of chemokine pathways (26, 27). Compartment-specific gene expression signatures were also identified, including an unexpected downregulation of cilium-associated genes in the nasal compartment. This finding suggests that innate defense mechanisms mediated by TLR signaling in the nose may require a reduction in mucociliary activity to enhance immune responsiveness. However, the functional implications of this observation remain to be fully elucidated and warrant further investigation.

The respiratory compartment most strongly modulated by flagellin was the bronchi, which exhibited the highest number of significantly enriched pathways and gene membership modules positively associated with flagellin treatment. Particularly those that influence granulocyte responses, with basal epithelial inflammatory mediators such as S100A8 and secretory epithelial mediators such as CXCL8 and CSF3R being significantly upregulated in response to flagellin in the bronchi. This is consistent with previous findings that flagellin mainly influences the human bronchial basal and secretory epithelial cells (43). A similar response has been observed in the pig. Flagellin triggers a strong inflammatory response in porcine bronchial epithelial cells differentiated under air-liquid culture, that is characterized by a transitory upregulation of *CXCL8, CXCL2* and *CCL20* (27).

Although most studies focus on the use of pigs as a model for human diseases, airway administration of flagellin also holds significant potential for porcine livestock. Mucosal exposure to flagellin has demonstrated protective activities in bacterial (27) and viral disease models, including influenza and post-influenza pneumococcal infection in the murine model (22). These protective effects are attributed to the pro-inflammatory activity on the airway epithelium and can provide a broad-spectrum defense against diverse pathogens. This characteristic of flagellin can particularly be advantageous for the porcine industry, as field respiratory diseases involve co-infections of virus and/or bacteria (44). Furthermore, the increased prevalence of antibiotic-resistant bacteria is a growing concern in porcine livestocks. For instance, *Actinobacillus pleuropneumoniae*, the major etiological bacterial agent of pleuropneumonia, has exhibited a significant increase over time in the resistance to several antibiotics (45). Studies in the murine model show that flagellin administration can enhance the efficacy of antibiotics against antibiotic-resistance bacteria without inducing detrimental inflammation (12, 16, 46). In a similar manner, our results show that flagellin administration to the airways produces a safe and transient pro-inflammatory lung environment. Further studies would be needed to determine whether flagellin can also enhance antibiotic efficacy in the pig model (47).

## Acknowledgements

We thank the Vaccine Development department of Statens Serum Institut (Denmark) for the manufacture and supply of FLAMOD, the skillful support by animal technicians from the Department Experimental Animals and Research of Wageningen Bioveterinary Research, and Florence Maurier for RNA-seq data submission.

## Funding

The study was funded by INSERM, Institut Pasteur de Lille, Université de Lille, the project FAIR and TRANSVAC2, that received funding from the European Union’s Horizon 2020 research and innovation program under grant agreement No 847786 and No 730964, respectively, and the VETBIONET European Union’s Horizon 2020 research infrastructures under grant agreement No 731014. The work was also supported by Agence Nationale de la Recherche (grant ANR-18-CE20-0024-01 to I.C.).

## Conflict-of-Interest Disclosure

JCS and NHV are the inventors of the patents WO2009156405, WO2011161491, and WO2015011254 that describes the use of FLAMOD as biologic against infectious diseases and the patent WO2023275292 on the formulation of FLAMOD. NHV is a cofounder of Cynbiose Respiratory. NHV was appointed as a scientific advisor (Novartis, European Commission, Immune Biosolution, ANR, Alvea/Telis Bioscience). NHV group has financial relationships with private-sector entities (Nemera, Aptar Pharma, Affilogic, 4P-Pharma, EuroAPI) outside the scope of the present work and Aerogen within the scope of this study. RMACL is named inventor on several vibrating mesh nebulizer patents. Authors declare no other competing interests.

## Author contributions

Contribution: MB, IC, NSZ and JCS planned studies. MB, LR, IC, DC, YLV, and NSZ performed experiments. RMACL and NHV optimized the aerosol delivery and supervised the aerosol administration to pigs, and revised the manuscript. DB, DH, CC, YZ and AGB performed RNA-seq and analysis. MB, IC, NSZ and JCS analyzed data and wrote the paper. NSZ and JCS supervised the project.

