## supplementary methods and figures and files for "Flagellin nebulization enhances respiratory immune responses in the porcine model": Supplemental_material_pig_nebulization_20250317.pdf

The supplementary materials include:

- Supplementary methods
- 4 supplementary figures
- 3 supplementary files (as separate files that list the differentially expressed genes, the associated biological processes using gene ontology, and the module identified by weighted gene co-expression network analysis, respectively)

### Supplementary methods

#### Weighted gene co-expression network analysis (WGCNA)

WGCNA was performed to find clusters of highly correlating genes across all transcriptomic samples of the respiratory tissues (1, 2). Using the normalized datasets as input data, the pairwise correlation coefficients were calculated between all pairs of genes using bi-weight mid-correlation (bicor) method, and were converted into an adjacency matrix using threshold power $\beta = 10$: the choice of $\beta$ is a value that correspond to an R-square of the scale-free fit greater than 0.8 and minimize the mean connectivity. The power value of 12 determines the scale-free topology of the resulting network. Signed topological overlap matrix was derived from the resulting adjacency matrix to measure the interconnectedness between genes. The co-expression network construction was performed under R WGCNA "blockwiseModule" function and the dynamic tree cut algorithm with height of 0.25, maxPOutliers of 0.05 and minimum module size of 30 genes. The expression profiles were summarized by the Module Eigengene (ME) representing the first principal components of each module. The module eigengenes were correlated with a binary matrix encoding the respiratory tissues. Gene Significance (GS) and Module membership (MM) were calculated using Pearson's correlation, to identify genes highly correlated with the compartments and those with the highest connectivity. As these two criteria help to identify key regulators within each module. Heatmap was generated to visualize the module-respiratory tissue correlation.

#### Gene expression analysis

Total RNA was reverse-transcribed with the High-Capacity cDNA Archive Kit (Applied Biosystems). cDNA was amplified using SYBR Green assays. Relative mRNA levels were determined by comparing the cycle thresholds (Ct) for the gene of interest and calibrator gene

*ACTB* (ΔCt) and were expressed as $2^{-\Delta\Delta Ct}$ values for flagellin-treated group compared to diluent-treated group. Ct upper limit was fixed to 35. Primer sequences are available upon request.

### Supplementary figures

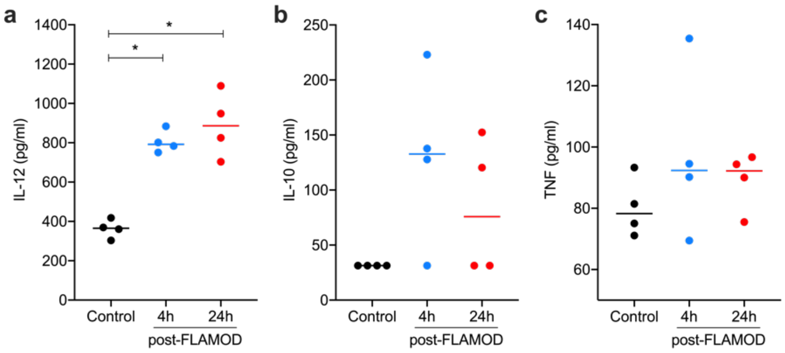


**Supplementary Figure 1. Nebulization of FLAMOD induces systemic immune responses.** Pigs were nebulized with diluent buffer (control group) or FLAMOD at 1.5 mg per pig. Animals were analyzed 4 h post-nebulization (4 h post-FLAMOD group) or 24 h post-nebulization (control and 24 h post-FLAMOD groups). Blood was sampled for preparation of serum and further analysis of production of cytokines IL-12 **(a)**, IL-10 **(b)** and TNF **(c)** was evaluated by ELISA. Each symbol represents one animal. Statistical analysis was performed using one-way ANOVA.

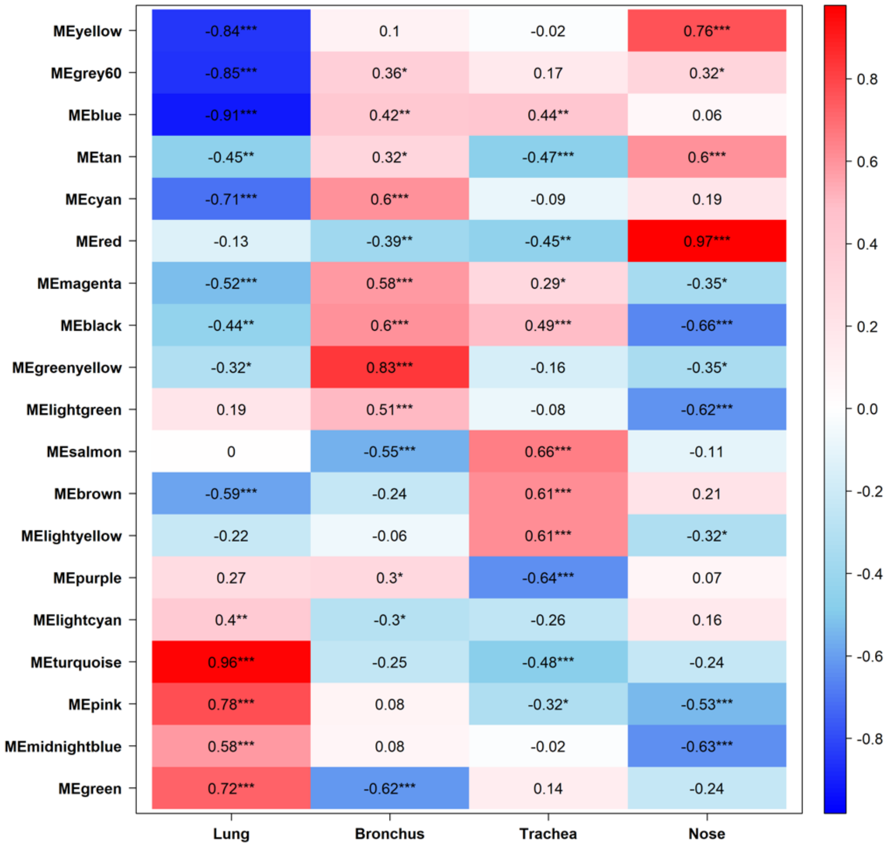


**Supplementary Figure 2. WGCNA analysis reveals that respiratory compartments show distinct patterns in response to FLAMOD nebulization.** Pigs were nebulized with diluent buffer (control group) or FLAMOD at 1.5 mg per pig. Animals were analyzed 4 h post-nebulization (4 h post-FLAMOD group) or 24 h post-nebulization (control and 24 h post-FLAMOD groups). Nasal mucosa (nose), trachea, primary bronchi, and lung samples were collected. Total RNA was extracted and subjected to RNA sequencing. Weighted gene co-expression network analysis (WGCNA) was performed on all samples (control, 4 h post-FLAMOD, and 24 h post-FLAMOD) from each tissue types. Gene significance ($p < 0.05$) and module membership (ME) is depicted per compartment, with each row referring to a module and each column to a respiratory compartment. Red represents positive correlation between network and compartment, while blue represents a negative correlation.

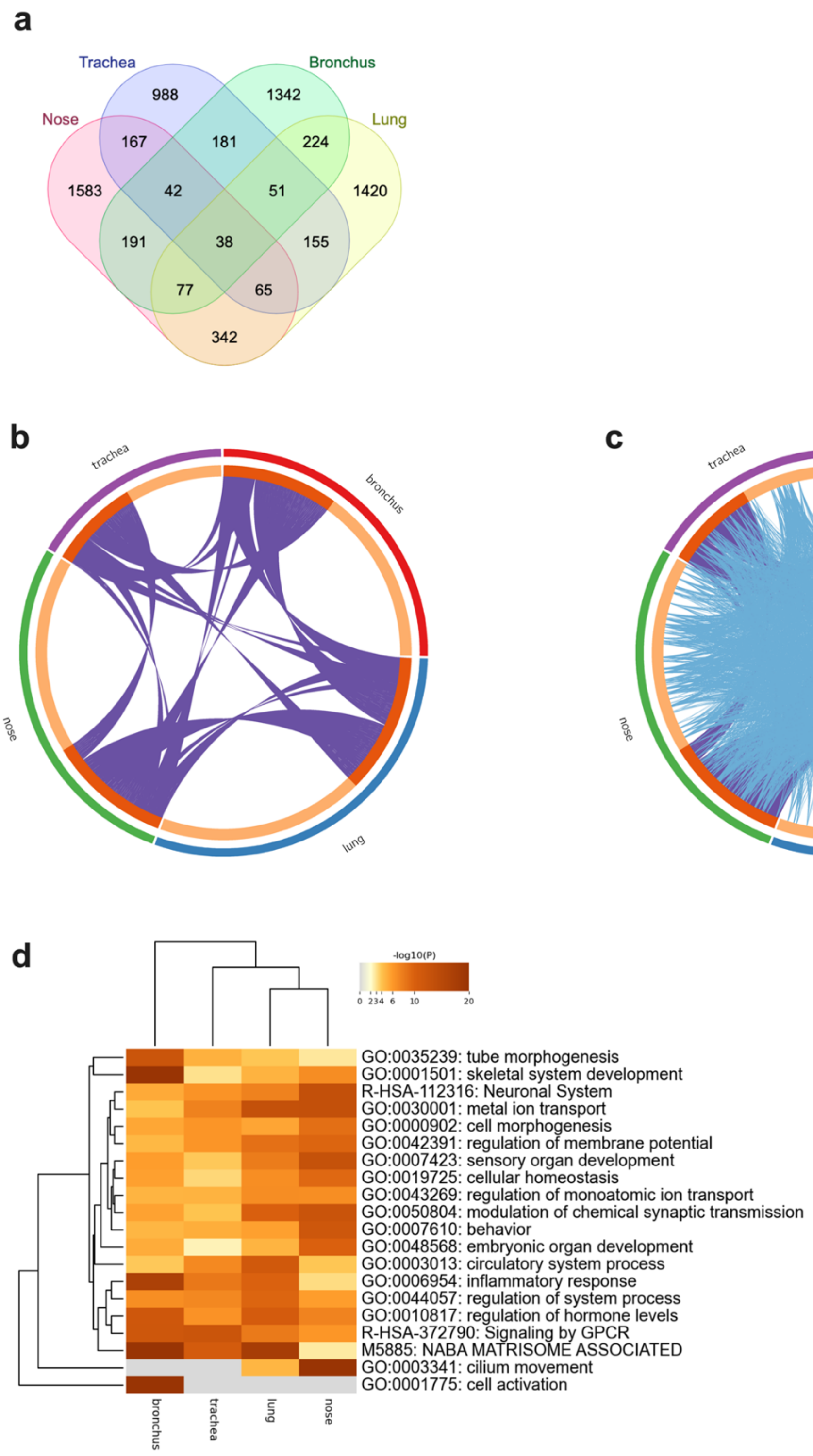


**Supplementary Figure 3. Flagellin aerosol delivery induces significant overlap in gene expression and biological pathways across respiratory compartments.** Pigs were nebulized with diluent buffer (control group) or FLAMOD at 1.5 mg per pig. Animals were analyzed 4 h post-nebulization (4 h post-FLAMOD group) or 24 h post-nebulization (control and 24 h post-FLAMOD groups). Nasal mucosa (nose), trachea, primary bronchi, and lung samples were collected. Total RNA was extracted and subjected to RNA sequencing. Gene expression overlap and pathway enrichment analyses were performed using Metascape (https://metascape.org). (a) Venn diagram showing the specific and overlapping differentially expressed genes across the different respiratory compartments. (b) Circos plot showing gene overlap between compartments. The outer arcs represent each compartment, while dark orange inner arcs denote genes shared between compartments, connected by purple lines. Light orange arcs represent genes unique to each compartment. (c) (c) Circos plot from (b) with additional functional pathway overlap analysis between compartments. Functional overlap is indicated by blue lines connecting genes sharing the same ontology term between compartments. (d) Dendogram of enrichment ontology clusters across compartments. The heatmap cells are colored by their p-values, white cells indicate the lack of enrichment for that term in the corresponding gene list.

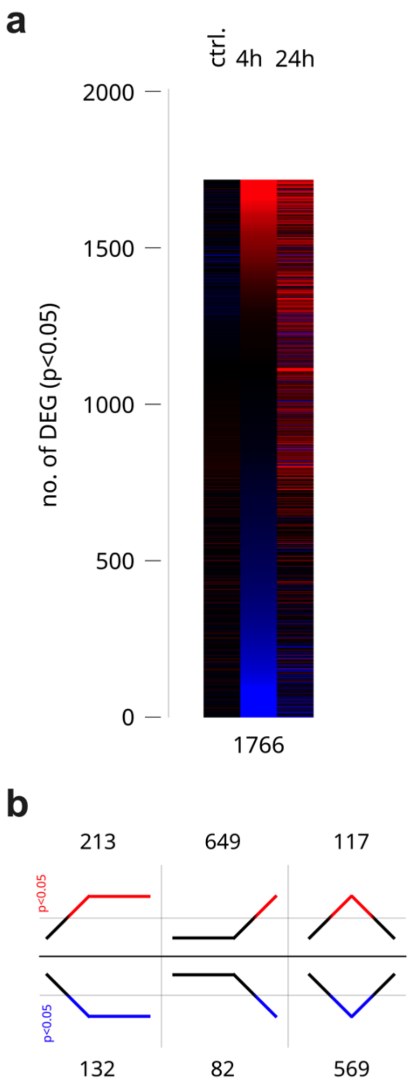


**Supplementary figure 4. FLAMOD nebulization patterns in the blood.** Pigs were nebulized with diluent buffer (control group) or FLAMOD at 1.5 mg per pig. Animals were analyzed 4 h post-nebulization (4 h post-FLAMOD group) or 24 h post-nebulization (control and 24 h post-FLAMOD groups). Blood samples were collected at 4 h post-nebulization (4h) or 24 h post-nebulization (24h). Total blood RNA was extracted and subjected to RNA sequencing **(a)** Heatmap illustrating the total number and relative expression change of regulated genes ($p<0.05$). **(b)** Time series analysis of each compartment showing the patterns of gene regulation as represented by six different line patterns, where the total number of genes statistically matching each pattern are indicated above or below the respective pattern.

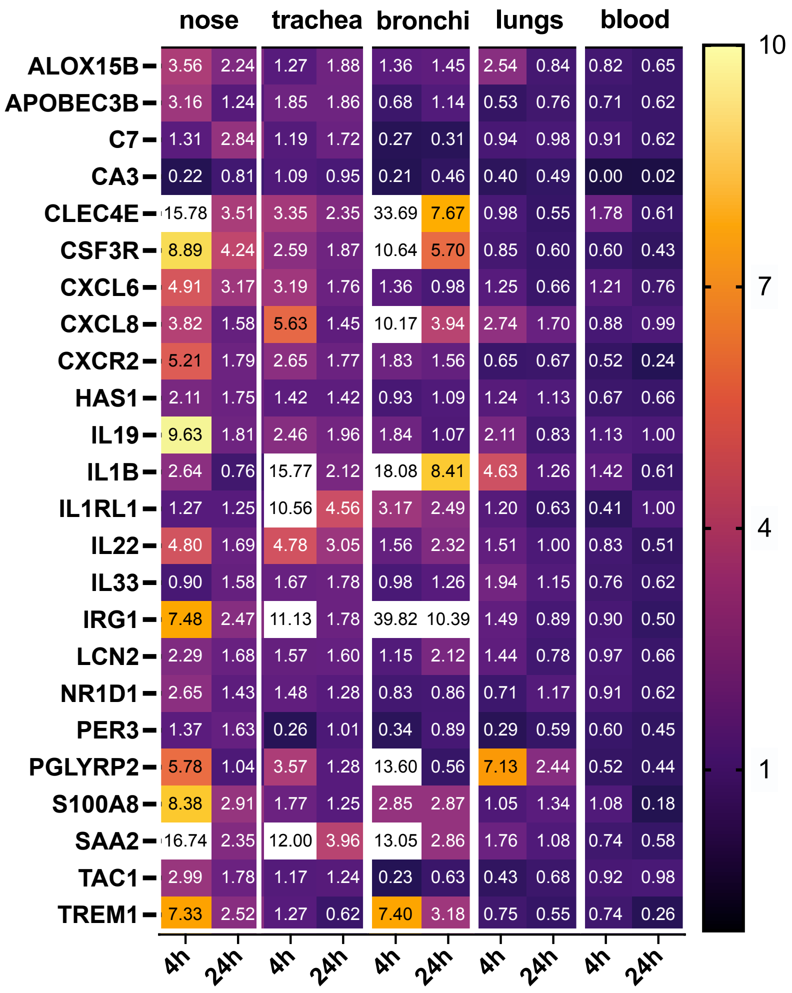


| | nose 4h | nose 24h | trachea 4h | trachea 24h | bronchi 4h | bronchi 24h | lungs 4h | lungs 24h | blood 4h | blood 24h |
|---|---|---|---|---|---|---|---|---|---|---|
| ALOX15B | 3.56 | 2.24 | 1.27 | 1.88 | 1.36 | 1.45 | 2.54 | 0.84 | 0.82 | 0.65 |
| APOBEC3B | 3.16 | 1.24 | 1.85 | 1.86 | 0.68 | 1.14 | 0.53 | 0.76 | 0.71 | 0.62 |
| C7 | 1.31 | 2.84 | 1.19 | 1.72 | 0.27 | 0.31 | 0.94 | 0.98 | 0.91 | 0.62 |
| CA3 | 0.22 | 0.81 | 1.09 | 0.95 | 0.21 | 0.46 | 0.40 | 0.49 | 0.00 | 0.02 |
| CLEC4E | 15.78 | 3.51 | 3.35 | 2.35 | 33.69 | 7.67 | 0.98 | 0.55 | 1.78 | 0.61 |
| CSF3R | 8.89 | 4.24 | 2.59 | 1.87 | 10.64 | 5.70 | 0.85 | 0.60 | 0.60 | 0.43 |
| CXCL6 | 4.91 | 3.17 | 3.19 | 1.76 | 1.36 | 0.98 | 1.25 | 0.66 | 1.21 | 0.76 |
| CXCL8 | 3.82 | 1.58 | 5.63 | 1.45 | 10.17 | 3.94 | 2.74 | 1.70 | 0.88 | 0.99 |
| CXCR2 | 5.21 | 1.79 | 2.65 | 1.77 | 1.83 | 1.56 | 0.65 | 0.67 | 0.52 | 0.24 |
| HAS1 | 2.11 | 1.75 | 1.42 | 1.42 | 0.93 | 1.09 | 1.24 | 1.13 | 0.67 | 0.66 |
| IL19 | 9.63 | 1.81 | 2.46 | 1.96 | 1.84 | 1.07 | 2.11 | 0.83 | 1.13 | 1.00 |
| IL1B | 2.64 | 0.76 | 15.77 | 2.12 | 18.08 | 8.41 | 4.63 | 1.26 | 1.42 | 0.61 |
| IL1RL1 | 1.27 | 1.25 | 10.56 | 4.56 | 3.17 | 2.49 | 1.20 | 0.63 | 0.41 | 1.00 |
| IL22 | 4.80 | 1.69 | 4.78 | 3.05 | 1.56 | 2.32 | 1.51 | 1.00 | 0.83 | 0.51 |
| IL33 | 0.90 | 1.58 | 1.67 | 1.78 | 0.98 | 1.26 | 1.94 | 1.15 | 0.76 | 0.62 |
| IRG1 | 7.48 | 2.47 | 11.13 | 1.78 | 39.82 | 10.39 | 1.49 | 0.89 | 0.90 | 0.50 |
| LCN2 | 2.29 | 1.68 | 1.57 | 1.60 | 1.15 | 2.12 | 1.44 | 0.78 | 0.97 | 0.66 |
| NR1D1 | 2.65 | 1.43 | 1.48 | 1.28 | 0.83 | 0.86 | 0.71 | 1.17 | 0.91 | 0.62 |
| PER3 | 1.37 | 1.63 | 0.26 | 1.01 | 0.34 | 0.89 | 0.29 | 0.59 | 0.60 | 0.45 |
| PGLYRP2 | 5.78 | 1.04 | 3.57 | 1.28 | 13.60 | 0.56 | 7.13 | 2.44 | 0.52 | 0.44 |
| S100A8 | 8.38 | 2.91 | 1.77 | 1.25 | 2.85 | 2.87 | 1.05 | 1.34 | 1.08 | 0.18 |
| SAA2 | 16.74 | 2.35 | 12.00 | 3.96 | 13.05 | 2.86 | 1.76 | 1.08 | 0.74 | 0.58 |
| TAC1 | 2.99 | 1.78 | 1.17 | 1.24 | 0.23 | 0.63 | 0.43 | 0.68 | 0.92 | 0.98 |
| TREM1 | 7.33 | 2.52 | 1.27 | 0.62 | 7.40 | 3.18 | 0.75 | 0.55 | 0.74 | 0.26 |

**Supplementary Figure 5. Immune gene expression is modulated by inhaled flagellin.** Pigs were nebulized with diluent buffer (control group) or FLAMOD at 1.5 mg per pig. Animals were analyzed and sampled 4 h post-nebulization (4 h post-FLAMOD group) or 24 h post-nebulization (control and 24 h post-FLAMOD groups). Samples were processed for RNA isolation and RT-qPCR analysis. Heatmaps displaying RT-qPCR data as fold changes in relative mRNA levels compared to the control group (normalized to 1). Upregulated gene expression (>1 fold change) is represented by a color gradient from purple to white, while downregulated expression (<1 fold change) is shown in a gradient from purple to black. Numeric annotations indicate the fold change for each gene in the respective tissue.
